# The SWELL1-LRRC8 complex regulates endothelial AKT-eNOS-mTOR signaling and vascular function

**DOI:** 10.1101/2020.08.04.236182

**Authors:** Ahmad F. Alghanem, Chau Ta, Joshua M. Maurer, Susheel K. Gunasekar, Ashutosh Kumar, Urooj Fatima, Chen Kang, Litao Xie, Oluwaseun Adeola, Javier Abello, Megan Riker, Macaulay Elliot-Hudson, Rachel A. Minerath, Amber Stratman, Chad E. Grueter, Robert F. Mullins, Rajan Sah

## Abstract

The endothelium responds to a multitude of chemical and mechanical factors in regulating vascular tone, angiogenesis, blood pressure and blood flow. The endothelial volume regulatory anion channel (VRAC) has been proposed to be mechano-sensitive, to activate in response to fluid flow/hydrostatic pressure and putatively regulate vascular reactivity and angiogenesis. Here, we show that the Leucine Rich Repeat Containing Protein 8a, LRRC8a (SWELL1) functionally encodes VRAC in human umbilical vein endothelial cells (HUVECs). Endothelial SWELL1 (SWELL1) expression positively regulates AKT-eNOS signaling while negatively regulating mTOR signaling, via a SWELL1-GRB2-Cav1-eNOS signaling complex. Endothelium-restricted SWELL1 KO (SWELL1 KO) mice exhibit enhanced tube formation from ex-vivo aortic ring explants in matrigel angiogenesis assays, develop hypertension in response to chronic angiotensin II infusion and have impaired retinal blood flow with both diffuse and focal blood vessel narrowing in the setting of Type 2 diabetes (T2D). These data demonstrate that SWELL1 antithetically regulates AKT-eNOS and mTOR signaling in endothelium and is required for maintaining vascular function, particularly in the setting of T2D.

## Introduction

The endothelium integrates mechanical and chemical stimuli to regulate vascular tone, angiogenesis, blood flow and blood pressure(1). Endothelial cells express a variety of mechanosensitive ion channels that regulate vascular function(2), including TRPV4(3-6) and Piezo1(7-9). The volume-regulated anion current (VRAC) is also prominent in endothelium, has been proposed to be mechano-sensitive(10), to activate in response to fluid flow/hydrostatic pressure(11) and putatively regulate vascular reactivity. However, the molecular identity of this endothelial ion channel has remained a mystery for nearly two decades.

*SWELL1* or *LRRC8a* (Leucine-Rich Repeat Containing Protein 8a) encodes a transmembrane protein first described as the site of a balanced translocation in an immunodeficient child with agammaglobulinemia and absent B-cells(12, 13). Subsequent work revealed the mechanism for this condition to be due to impaired SWELL1-dependent GRB2-PI3K-AKT signaling in lymphocytes, resulting in a developmental block in lymphocyte differentiation(14). Thus, for ∼11 years, SWELL1 was conceived of as a membrane protein that regulates PI3K-AKT mediated lymphocyte function(12, 13). Although SWELL1 had been predicted to form a hetero-hexameric ion channel complex with other LRRC8 family members(15), it was not until 2014 that SWELL1/LRRC8a was shown to form an essential component of the volume-regulated anion channel (VRAC)(16, 17), forming hetero-hexamers with LRRC8b-e(17, 18). Therefore, historically, SWELL1-LRRC8 complex was first described as a membrane protein that signaled via protein-protein interactions and then later found to form an ion channel signaling complex.

We showed previously that SWELL1 (LRRC8a) is an essential component of VRAC in adipocytes that is required for insulin-PI3K-AKT2 signaling to mediate adipocyte hypertrophy and systemic glucose homeostasis(19-21). The PI3K-AKT-eNOS signaling pathway is central to transducing both mechanical stretch(22) and hormonal inputs (insulin) to regulate endothelial nitric oxide synthase (eNOS) expression and activity, which, in turn regulates vasodilation (blood flow and pressure), inhibits leukocyte aggregration, and limits proliferation of vascular smooth muscle cells (atherosclerosis). Indeed, insulin resistance is thought to be a systemic disorder in the setting of Type 2 diabetes (T2D), affecting endothelium in addition to traditional metabolically important tissues, such as adipose, liver, and skeletal muscle(23-25). In fact, insulin resistant endothelium and the resultant impairment in PI3K-AKT-eNOS signaling has been proposed to underlie much of the endothelial dysfunction observed in the setting of obesity and T2D, predisposing to hypertension, atherosclerosis and vascular disease(23-25).

In this study, we demonstrate that VRAC is SWELL1-dependent in endothelium, associates with GRB2, caveolin-1 (Cav1), endothelial nitric oxide synthase (eNOS), and regulates PI3K-AKT-eNOS, ERK1/2 and mTOR signaling – suggesting that SWELL1-LRRC8 channel complexes link insulin and mechano-signaling in endothelium. SWELL1-dependent AKT-eNOS, ERK1/2 and mTOR signaling influences angiogenesis, blood pressure and vascular function *in vivo*, while impaired endothelial SWELL1-LRRC8 signaling predisposes to vascular dysfunction in the setting of diet-induced T2D.

## Results

### SWELL1 functionally encodes VRAC in endothelium

The volume-regulatory anion current (VRAC) has been measured and characterized in endothelial cells for decades but the molecular identity of this endothelial ion channel remains elusive(10, 11, 26). To determine if the leucine-rich repeat containing membrane protein SWELL1 (LRRC8a) recently identified in cell lines (16, 17) is required for VRAC in endothelial cells, as it is in adipocytes (19), pancreatic β-cells (27, 28), nodose neurons (29) and spermatozoa (30), we first confirmed robust SWELL1 protein expression by Western blot (**Figure 1A)** and immunostaining (**Figure 1B**) in human umbilical vein endothelial cells (HUVECs). SWELL1 protein expression is substantially reduced upon adenoviral transduction with a short-hairpin RNA directed to SWELL1 (Ad-shSWELL1-mCherry) as compared to a scrambled control (Ad-shSCR-mCherry). Next, we measured hypotonically-induced (210 mOsm) endothelial VRAC currents in HUVECs. These classic outwardly rectifying hypotonically-induced VRAC currents are prominent in HUVECs, largely blocked by the VRAC inhibitor 4-(2-Butyl-6,7-dichloro-2-cyclopentyl-indan-1-on-5-yl) oxobutyric acid (DCPIB; **Figure 1C&D**), and significantly suppressed upon shSWELL1-mediated SWELL1 knock-down (**Figure 1E&F**), consistent with SWELL1 functionally encoding endothelial VRAC.

**Figure 1.**
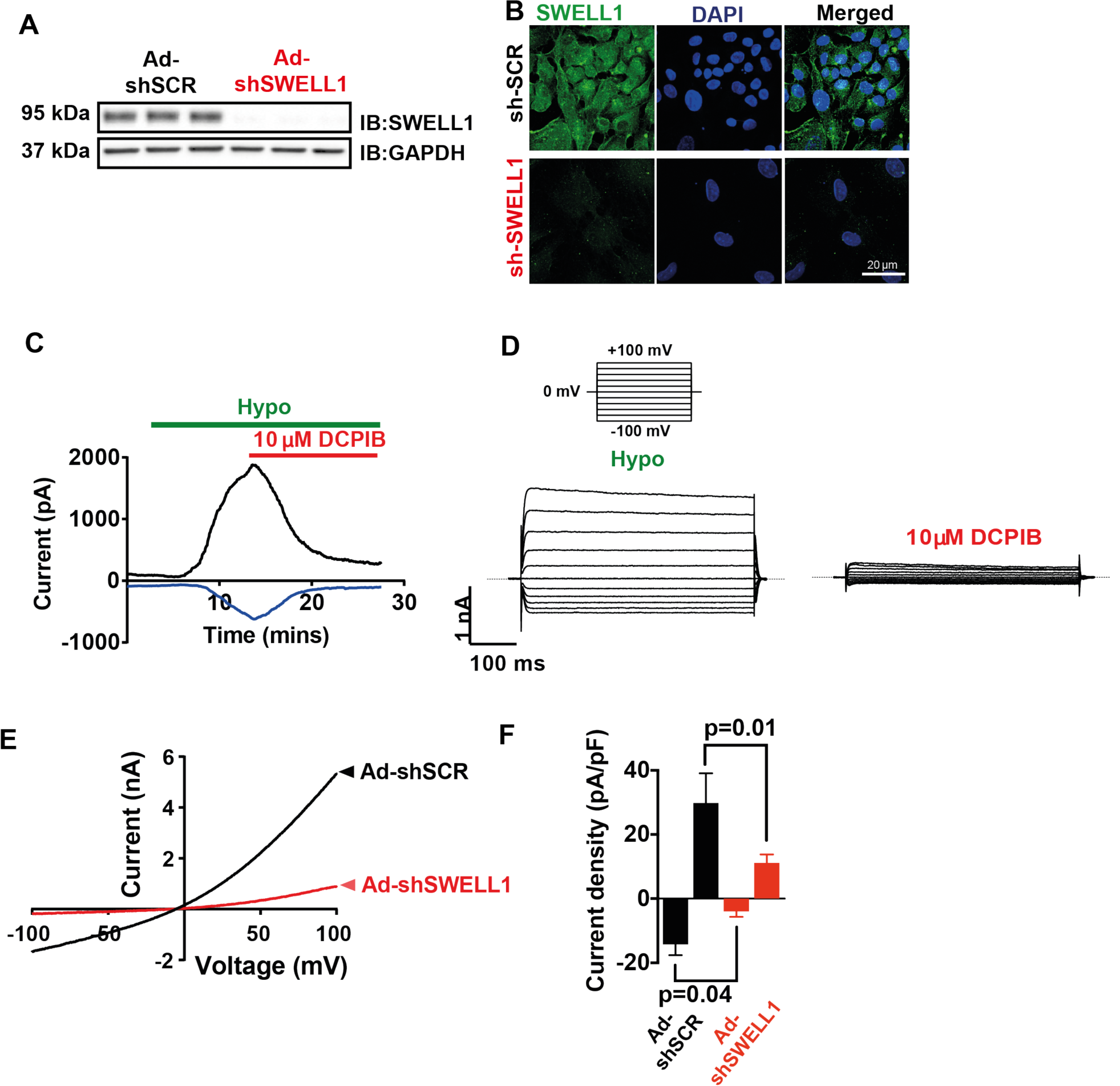
SWELL1 mediates VRAC currents in human umbilical vein endothelial cells (HUVECs). **A**, SWELL1 western blot in HUVECs transduced with adenovirus expressing a short hairpin RNA directed to SWELL1 (Ad-shSWELL1) compared to control scrambled short hairpin RNA (Ad-shSCR). GAPDH is used as loading control. **B**, Immunofluorescence staining of the HUVECs transduced with Ad-shSWELL1 and Ad-shSCR. **C**, Current-time relationship of VRAC (hypotonic, 210 mOsm) in Ad-shSCR transduced HUVEC and co-application of 10 µM DCPIB. **D**, Representative current traces upon hypotonic activation (left) during voltage steps (from -100 to +100 mV, shown in inset) and inhibition by DCPIB (right). **E**, Current-voltage relationship of VRAC during voltage ramps from -100 mV to +100 mV after hypotonic swelling in HUVECs transduced with Ad-shSCR and Ad-shSWELL1. **F**. Mean current outward and inward densities at +100 and -100 mV (n,_sh-SCR_=4 cells; n,_shSWELL1_=6 cells). Data are shown as mean ± s.e.m. *p <0.05; **p <0.01; unpaired t-test for **F**.

### SWELL1 regulates PI3K-AKT-eNOS, ERK and mTOR signaling in endothelium

Previous studies in adipocytes demonstrate that SWELL1 regulates insulin-PI3K-AKT signaling, adipocyte expansion and systemic glycemia, whereby SWELL1 loss-of-function induces an insulin-resistant pre-diabetic state (19, 20). Insulin signaling is also important in regulating endothelium and vascular function (24, 25, 31). Moreover, insulin-resistance in Type 2 diabetes (T2D) is considered a systemic disorder and insulin-resistant endothelium is postulated to underlie impaired vascular function in T2D (24, 25). As SWELL1 is highly expressed in endothelium (**Figure 1**), and PI3K-AKT-eNOS signaling critical for endothelium-dependent vascular function(32), we next examined AKT-eNOS, ERK1/2 and mTOR signaling in SWELL1 KD compared to control HUVECs under basal conditions. Basal phosphorylated AKT2 (pAKT2, **Figure 2A&D**), pAKT1 (**Figure 2A&E**), p-eNOS (**Figure 2A&C**), pERK1/2 (**Figure 2A&F**) are abrogated in HUVECs upon SWELL1 KD, indicating that SWELL1 contributes to AKT-eNOS, and ERK signaling in endothelium. Curiously, basal pS6 ribosomal protein, indicative of mTOR signaling, is augmented in SWELL1 KD HUVECs compared to control (**Figure 2A&G**), suggesting SWELL1 to be a negative regulator of mTOR in endothelium. As a complementary approach, we used siRNA mediated SWELL1 knock-down using a silencer select siRNA targeting SWELL1 mRNA with a different sequence from shSWELL1. siRNA mediated SWELL1 KD in HUVECs yielded nearly identical results to the shRNA KD approach (**Figure 2-Figure Supplement 1**). In summary, SWELL1 expression level regulates AKT, eNOS, ERK and mTOR signaling in endothelium.

**Figure 2.**
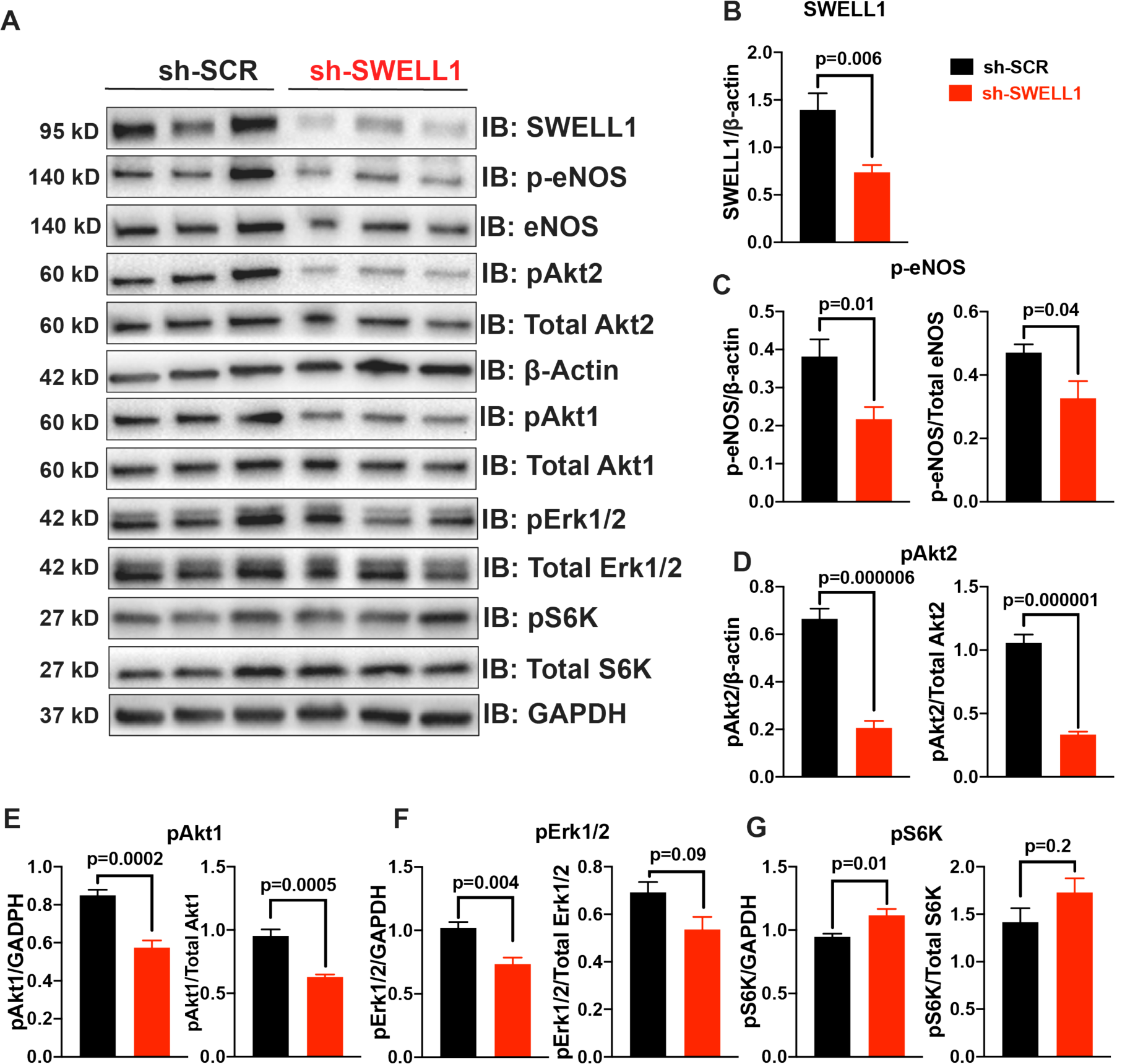
SWELL1 regulates PI3K-AKT-eNOS, ERK and mTOR signaling in endothelium. (**A**), Western blots of SWELL1, pAkt2, pAkt1, Akt2, Akt1, pErk1/2, Erk1/2, p-eNOS, eNOS, pS6K ribosomal protein, S6K ribosomal protein, GAPDH, and β-Actin in Ad-shSCR and Ad-shSWELL1 transduced HUVECS under basal conditions. Quantification of SWELL1/ β-Actin (**B**), p-eNOS/ β-actin, p-eNOS/Total eNOS (**C**), pAkt2/ β-actin, pAkt2/Total Akt2 (**D**), pAkt1/GAPDH, pAkt1/Total Akt1 (**E**), pERK1/2 /GAPDH, pErk1/2 /Total Erk1/2 (**F**), pS6 ribosomal protein/GAPDH, and pS6K ribosomal protein/Total S6K ribosomal protein (**G**). N=6 independent experiments. Significance between the indicated groups in all blots were calculated using a two-tailed Student’s t-test. P-values are illustrated on figures. Error bars represent mean ± s.e.m.

**Figure 2- Figure Supplement 1.**
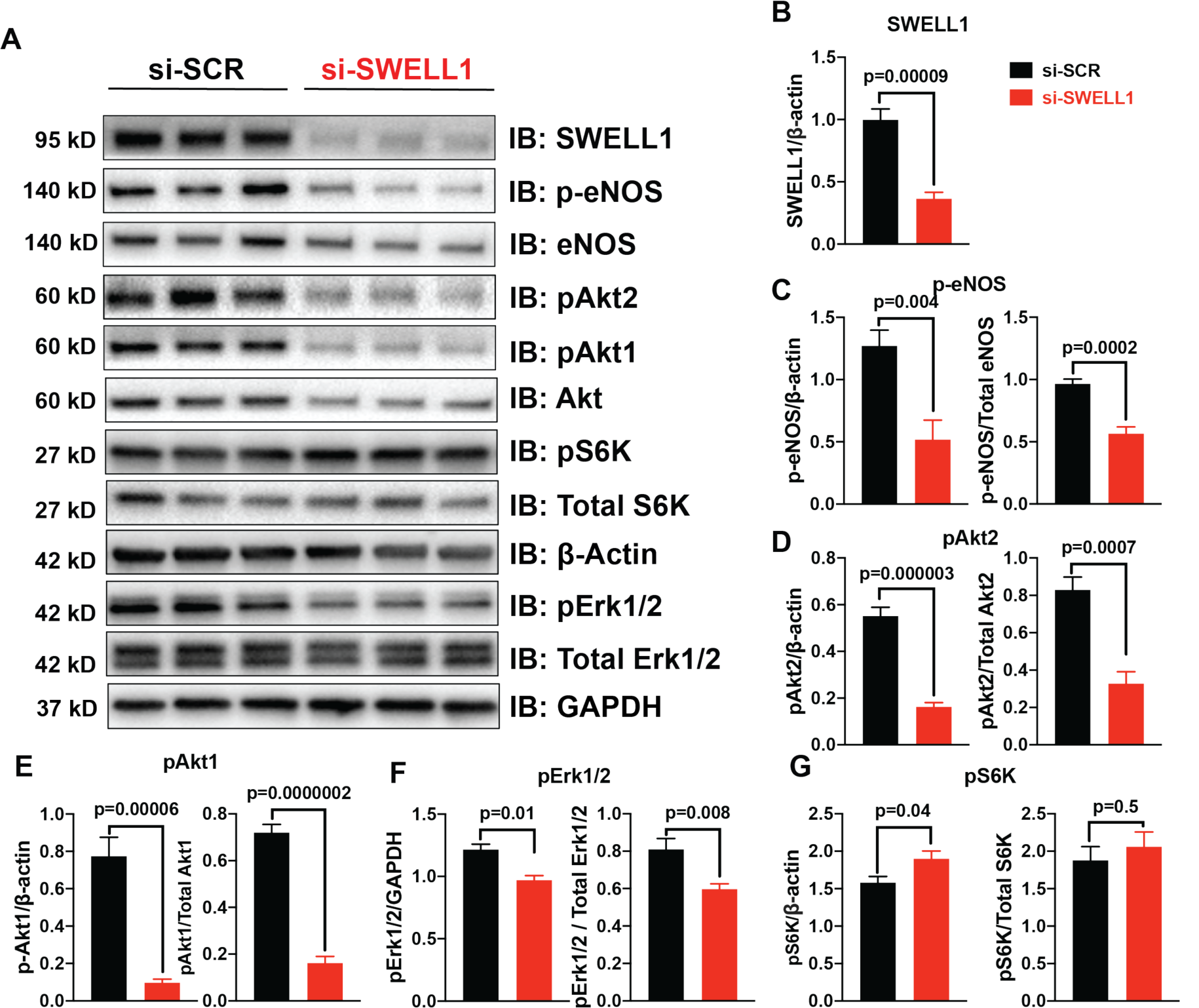
SWELL1 regulates PI3K-AKT-eNOS, ERK and mTOR signaling in endothelium. (**A**), Western blots of SWELL1, pAkt2, pAkt1, Akt, pErk1/2, Erk1/2, p-eNOS, eNOS, pS6K ribosomal protein, S6K ribosomal protein, and β-Actin in si-SCR and si-SWELL1 transduced HUVECS under basal conditions. Quantification of SWELL1/ β-Actin (**B**), p-eNOS/ β-actin, p-eNOS/Total eNOS (**C**), pAkt2/ β-actin, pAkt2/Total Akt (**D**), pAkt1/ β-actin, pAkt1/Total Akt (**E**), pERK1/2 / β-actin, pErk1/2 /Total Erk1/2 (**F**), pS6 ribosomal protein/ β-actin, and pS6K ribosomal protein/Total S6K ribosomal protein (**G**). N=6 independent experiments. Significance between the indicated groups in all blots were calculated using a two-tailed Student’s t-test. P-values are illustrated on figures. Error bars represent mean ± s.e.m.

### SWELL1 interacts with GRB2, Cav1 and eNOS and mediates stretch-dependent eNOS signaling

In adipocytes, the mechanism of SWELL1-mediated regulation of PI3K-Akt signaling involves SWELL1/GRB2/Cav1 molecular interactions (19). To determine if SWELL1 resides in a similar macromolecular signaling complex in endothelium we immunoprecipitated (IP) endogenous GRB2 from HUVECs. Upon GRB2 IP, we detect SWELL1 protein in shSCR treated HUVECs and less SWELL1 upon GRB2 IP from shSWELL1-treated HUVECs, consistent with a SWELL1-GRB2 interaction (**Figure 3A&B**). In addition, with GRB2 IP we also detect both Cav1 (**Figure 3A&B**) and eNOS (**Figure 3B**). These data suggest that endothelial SWELL1 resides in a signaling complex that includes GRB2, Cav1 and eNOS, consistent with the findings that GRB2 and Cav1 interact, and that Cav1 regulates eNOS via a direct interaction (33-35). Also, GRB2 has been shown to regulate endothelial ERK, AKT and JNK signaling (36). Moreover, these data are also in-line with the notion that caveoli form mechanosensitive microdomains (37-39) that regulate VRAC (40, 41) and that VRAC can be activated by mechanical stimuli in a number of cell types, including endothelium (10, 26, 42-45).

**Figure 3.**
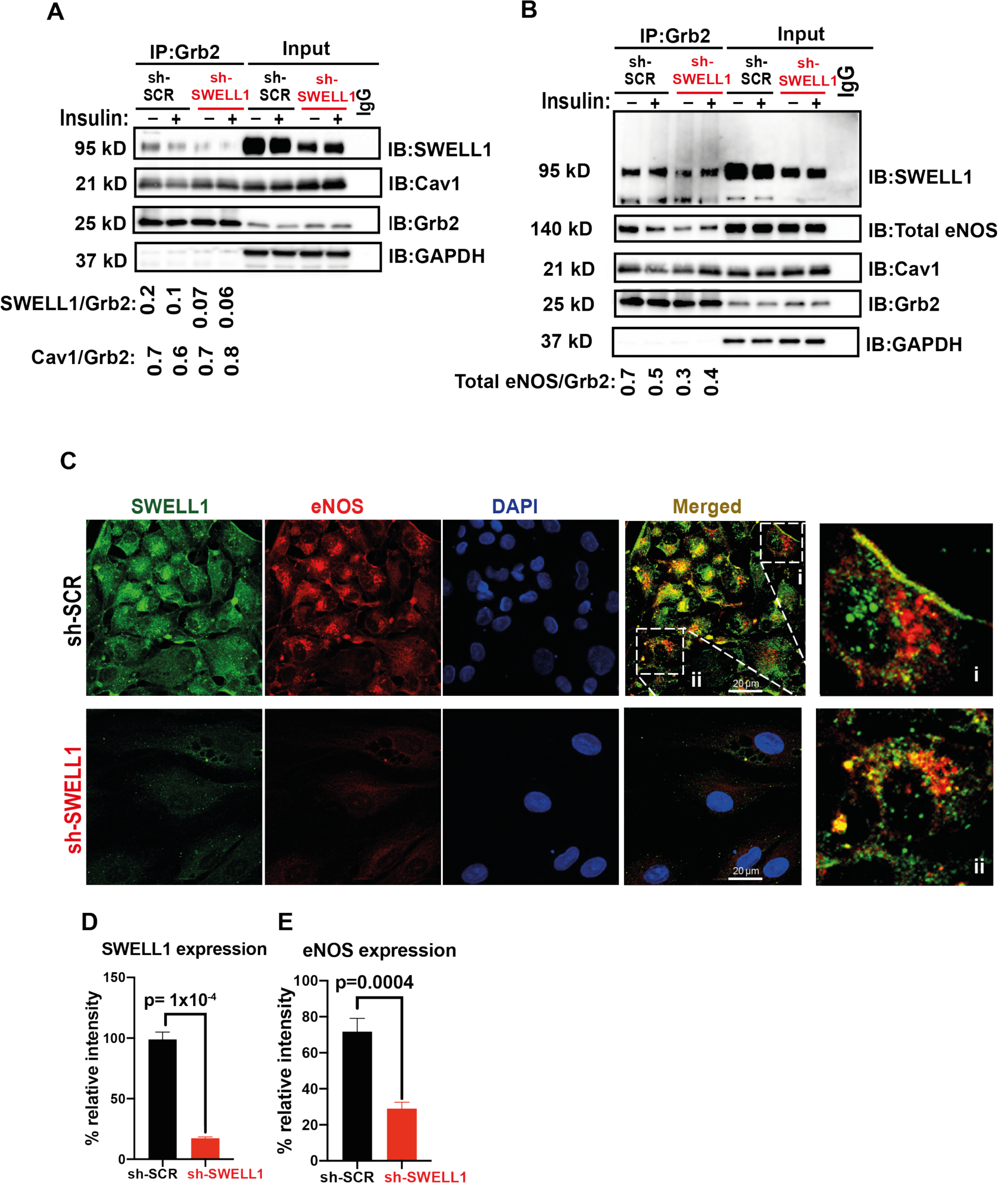
SWELL1 interacts with Grb2, Cav1 and eNOS in human endothelium. **A**, GRB2 immunoprecipitation from Ad-shSCR and Ad-shSWELL1 transduced HUVECs and immunoblot with SWELL1, Cav1 and GRB2 antibodies. Densitometry values for GRB2 co-immunoprecipitated SWELL1 (SWELL1/GRB2) and GRB2 co-immunoprecipitated Cav1 (Cav1/GRB2). GAPDH serves as loading control for input samples. (**B**) GRB2 immunoprecipitation from Ad-shSCR and Ad-shSWELL1 transduced HUVECs and immunoblot with SWELL1, eNOS, Cav1 and GRB2 antibodies. Insulin-stimulation with 100 nM insulin for 10 minutes. Densitometry values for GRB2 co-immunoprecipitated eNOS (eNOS/GRB2). Representative blots from 3 independent experiments. (**C**) Representative endogenous SWELL1 and eNOS immunofluorescence staining in Ad-shSCR and Ad-shSWELL1 transduced HUVECs. Representative image from 6 independent experiments. (**D-E**) Quantification of SWELL1 (**D**, n = 6) and eNOS (**E**, n = 6) immunofluorescence staining upon SWELL1 KD. Evidence of SWELL1-eNOS colocalization (C, insets) in plasma membrane (**i**) and perinuclear regions (**ii**). Significance between the indicated groups are calculated using a two-tailed Student’s t-test. P-values are indicated on figures. Error bars represent mean ± s.e.m.

We also examined the relationship between SWELL1 and eNOS protein expression and localization in HUVECs by immunofluorescence (IF) staining (**Figure 3C**). Similar to observed by Western blot (**Figure 2A, and Figure 2-Figure Supplement 1**), IF staining reveals that reductions in SWELL1 expression correlate with reduced eNOS expression (**Figure 3C-E, Figure 3-Figure Supplement 1**). Moreover, SWELL1 and eNOS co-localize in plasma membrane and peri-nuclear intracellular domains (**Figure 3C, inset**), consistent with the IP data revealing a SWELL1-GRB2-Cav-eNOS interaction (**Figure 3A&B**), and also with previously described intracellular eNOS localization(46).

**Figure 3 - Figure Supplement 1.**
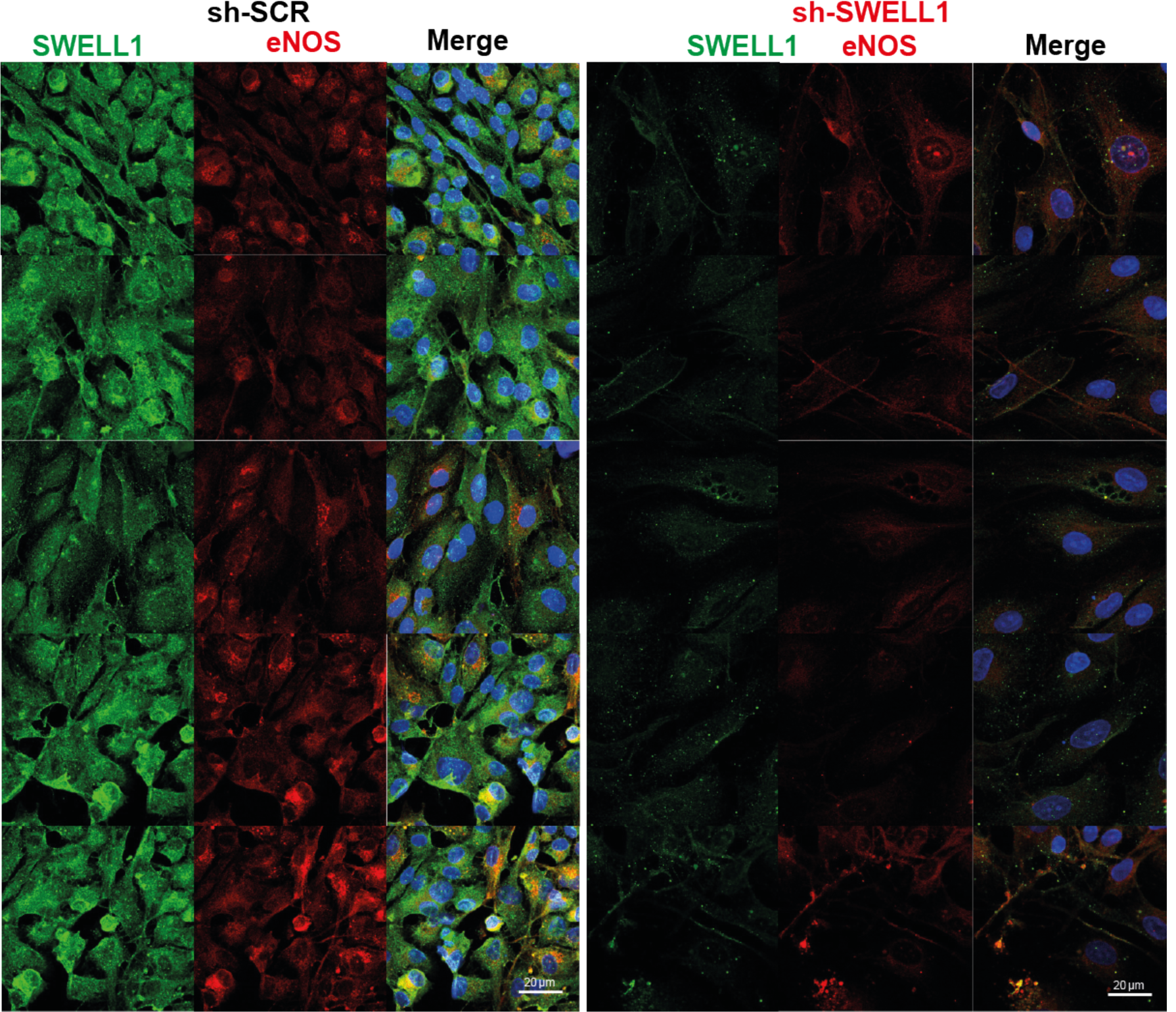
SWELL1 co-localizes with eNOS and regulates eNOS expression. SWELL1 (green) and eNOS (red) immunofluorescence staining of HUVEC transduced with either Ad-shSCR or Ad-shSWELL1. Scale bar is 20 µm. DAPI (blue) labels nuclei.

Given that endothelial cells respond to stretch stimuli to regulate vascular tone via activation of eNOS, we next examined the SWELL1-dependence of stretch-induced AKT, ERK1/2 and eNOS signaling in HUVECs (**Figure 4**). Stretch (5%) is sufficient to stimulate AKT1 and AKT2 signaling (**Figure 4A-C**), though not ERK1/2 signaling (**Figure 4D**) in HUVECs, and all are blunted in SWELL1 KD HUVECS (**Figure 4A-D**). Similarly, we observe abrogation of time-dependent p-eNOS signaling with 5% stretch in SWELL1 KD HUVECS compared to control (**Figure 4E&F**). Taken together, these data position SWELL1 as a regulator stretch-mediated PI3K-AKT-eNOS signaling in endothelium via a SWELL1-GRB2-Cav1-eNOS signaling complex.

**Figure 4.**
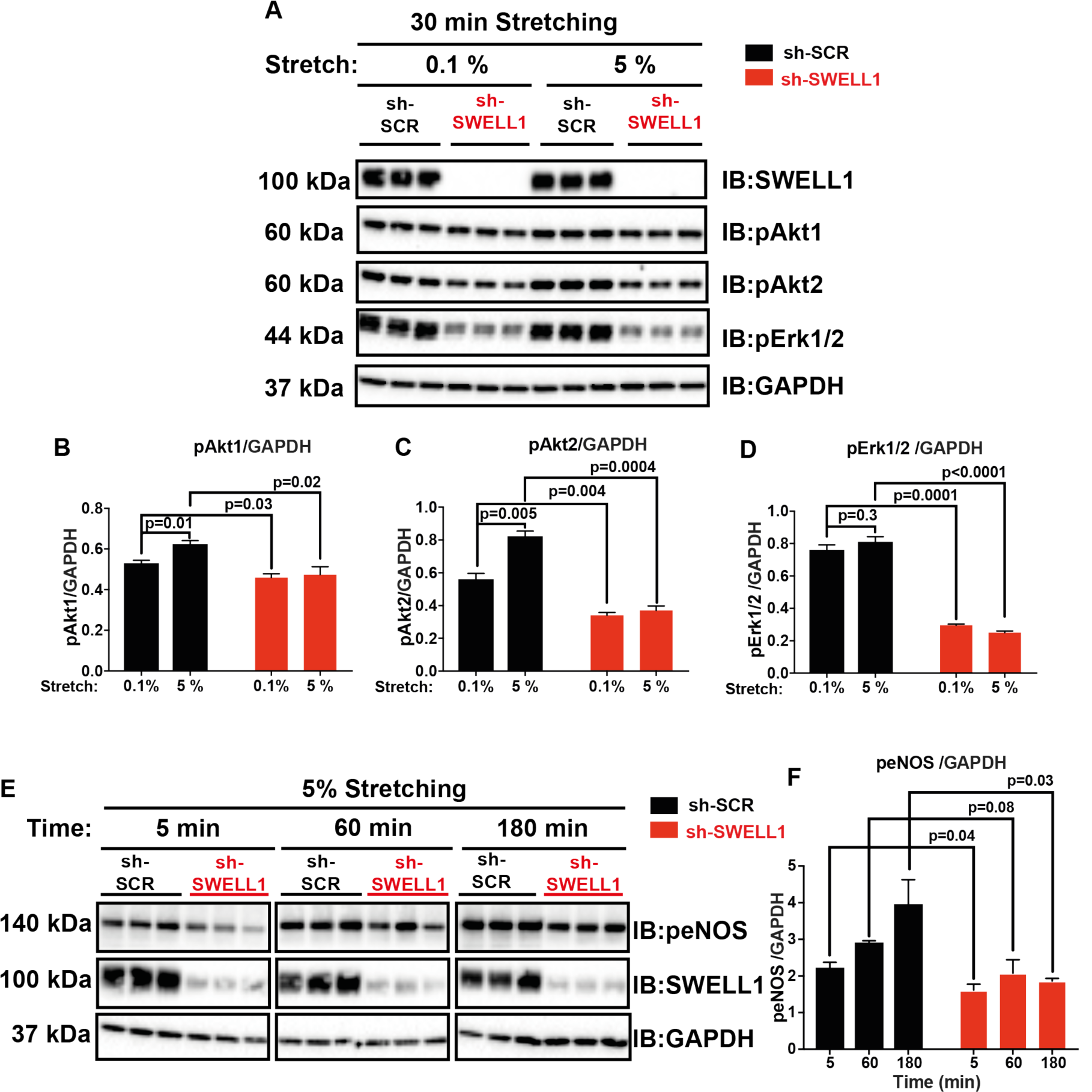
SWELL1 is required for intact stretch-induced AKT-eNOS signalling. **A**, Western blot of SWELL1, pAKT1, pAKT2, pERK1/2 in response to 30 minutes of 0.1% and 5% static stretch in Ad-shSCR and Ad-shSWELL1 transduced HUVECs. GAPDH is used as a loading control. (**B**-**D**) Densitometry quantification from A of pAKT1 (**B**), pAKT2 (**C**) and pErk1/2 (**D**). (**E**) Western blot of peNOS, SWELL1, in response to 5% static stretching for 5, 60 and 180 min in Ad-shSCR and Ad-shSWELL1 transduced HUVECs. (**F**) Densitometry quantification from E of eNOS. GAPDH is used as a loading control. Significance between the indicated groups in all blots were calculated using a two-tailed Student’s t-test. P-values are illustrated on figures. Error bars represent mean ± s.e.m.

To examine the functional consequences of endothelial SWELL1 ablation *in vivo* we generated endothelial-targeted SWELL1 KO mice (eSWELL1 KO) by crossing SWELL1 floxed mice (19, 27) with the endothelium-restricted CDH5-Cre mouse (*CDH5-Cre;SWELL1*^*fl/fl*^; **Figure 5A**).

**Figure 5.**
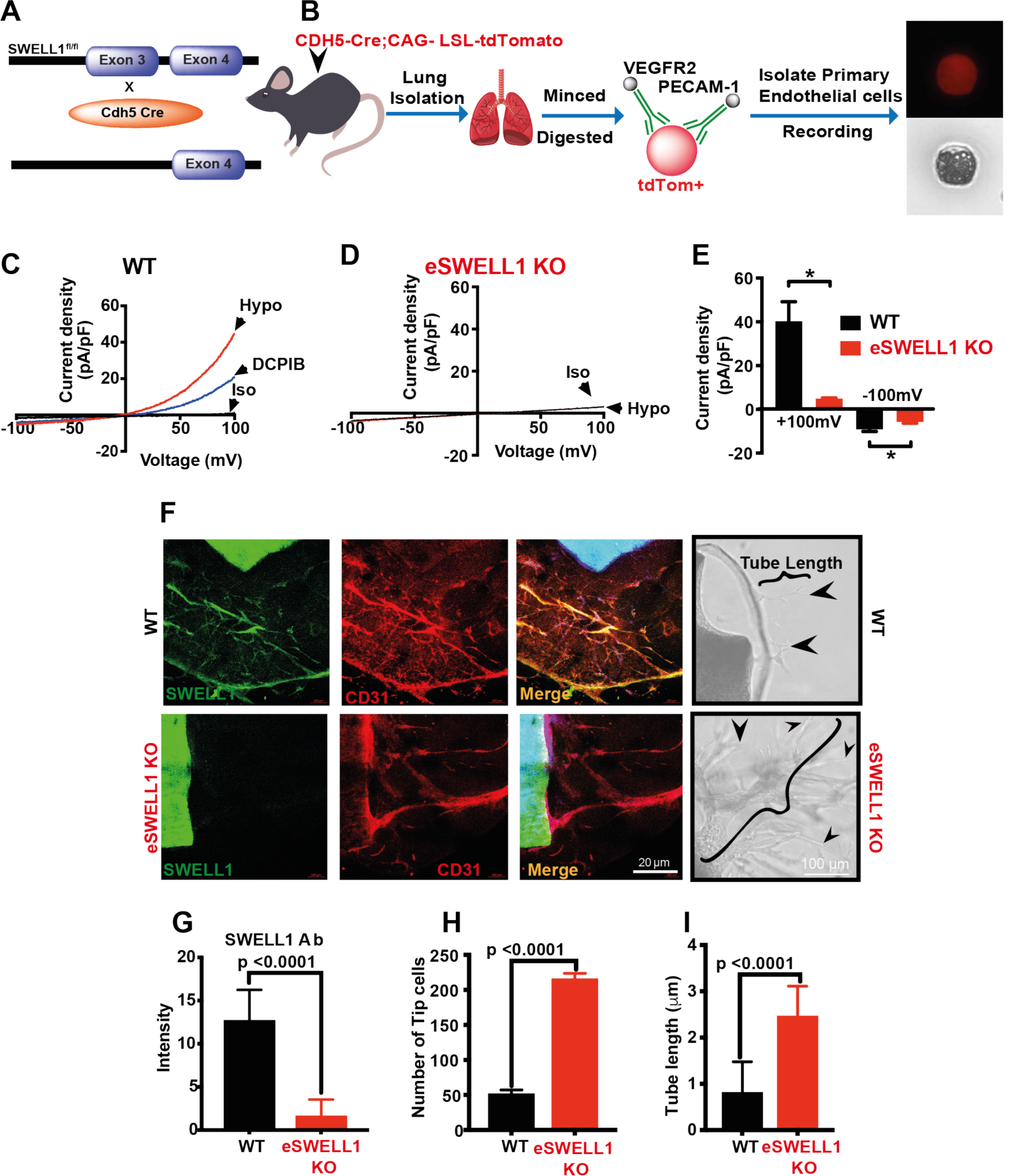
Endothelium-specific SWELL1 KO mice exhibit enhanced tube formation from aortic explants. **A**, Strategy for endothelium targeted SWELL1 ablation to generate eSWELL1 KO. (**B**) Isolation of murine primary endothelial cells from WT and eSWELL1 KO using tdTomato reporter mice. **C-D**, Current-voltage relationships of VRAC in isotonic (Iso, 300 mOsM) and hypotonic (Hypo, 210 mOsm) solution in response to voltage ramps from -100 to +100 mV over 500 ms in WT (**C**) and KO (**D**) primary murine endothelial cells. DCPIB (10 *μ*M) inhibition in C (WT). (E) Mean outward (+100mV) and inward (−100 mV) currents from WT (n=3 cells) and eSWELL1 KO (n = 3 cells). (**F**) Ex-vivo aorta sprouting assay performed in aortic rings isolated from WT and eSWELL1 KO mice and cultured in FGM media for 3 days at 37oC. Immunofluoresence staining with antibodies to SWELL1 (green), CD31 (red), SWELL1+CD31+ (Merge) and bright field images show endothelial cell tubes sprouting from WT and eSWELL1 KO aortic rings (black arrow heads). (**G-I**) Quantification of SWELL1 immunofluorescence signal (G, WT = 15, KO = 15), number of tip cells (**H**, WT = 26, KO = 31), and endothelial tube length (**I**, WT = 30, KO = 30) in WT and eSWELL1 KO aortic explants. Statistical significance between the indicated values calculated using a two-tailed Student’s t-test. P-values are illustrated on figures. Error bars represent mean ± s.e.m. n = 3, independent experiments.

Patch-clamp recordings of primary endothelial cells isolated from WT and eSWELL1 KO mice (**Figure 5B**) revealed robust hypotonically-activated currents (Hypo, 210 mOsm) in WT endothelial cells, that are DCPIB inhibited (**Figure 5C&E**), while eSWELL1 KO endothelial cells exhibit markedly reduced hypotonically-activated currents (**Figure 5D&E**). Immunofluorescence staining of aortic ring explants revealed that SWELL1 ablation from CD31+ primary endothelial cells significantly enhanced *ex-vivo* sprouting angiogenesis from these explants (**Figure 5F&G**), based on both tube length and number of tip cells in eSWELL1 KO mice as compared to WT mice (**Figure 5&H-I**), suggesting that SWELL1 regulates angiogenesis. Indeed, genome-wide transcriptome analysis of SWELL1 KD HUVEC compared to control (RNA sequencing) reveal multiple pathways enriched regulating angiogenesis, migration and tumorigenesis, including GADD45, IL-8, p70S6K (mTOR), TREM1, angiopoeitin and HGF signaling (**Figure 6, Supplementary Table 1&2**). Also, notable are statistically significant increases in VEGFA (1.6-fold) and CD31 (2.0-fold) expression in SWELL1 KD HUVECs, both of which are pro-angiogenic and associated with mTORC1 hyperactivation(47). Pathways linked to cell adhesion and renin-angiotensin signaling are also enriched - both pathways and processes that are known to be altered in vasculature in the setting of atherosclerosis and Type 2 diabetes (T2D). Finally, the trends toward reduced eNOS protein expression observed upon SWELL1 KD in HUVECs (**Figure 2A, Supplementary Fig 1A, Fig. 3C, Figure 3-Figure Supplement 1**) are associated with reduced eNOS mRNA expression (0.54-fold in SWELL1 KD HUVEC, p < 10^−4^).

**Figure 6.**
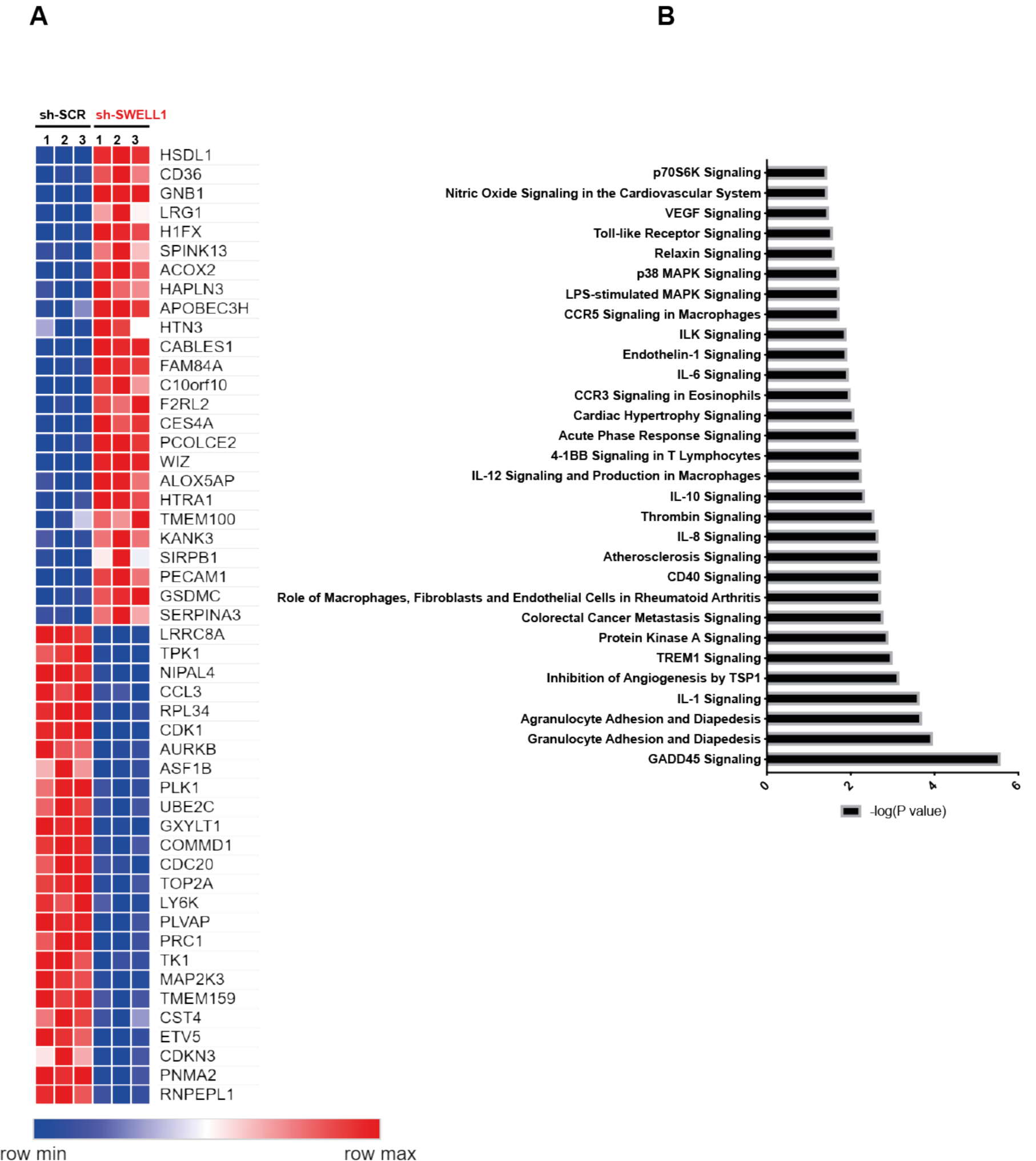
RNA sequencing of Ad-shSCR and Ad-shSWELL1 transduced HUVECs. **A**, Heatmap analysis displaying top 25 upregulated or 25 downregulated genes between shSCR and shSWELL1. (**B**) IPA canonical pathway analysis of genes significantly regulated by shSWELL1 in comparison to shSCR. n = 3 for each group. For analysis with IPA, FPKM cutoffs of 1.5, fold change of ≥1.5, and false discovery rate < 0.05 were utilized for significantly differentially regulated genes.

### eSWELL1 KO mice exhibit mild angiotensin-II stimulated hypertension and impaired retinal blood flow in the setting of Type 2 diabetes

Based on our findings that SWELL1 regulates AKT-eNOS signaling in endothelium, and that eNOS signaling is central to blood pressure regulation, we next examined blood pressures in eSWELL1 KO mice compared to WT controls (SWELL1^fl/fl^ mice). Male mice exhibit no significant differences in systolic blood pressure under basal conditions (**Figure 7A**), while female mice are mildly hypertensive relative to WT mice (**Figure 7B**). However, after 4 weeks of angiotensin-II infusion (Ang II), male eSWELL1 KO mice develop exacerbated systolic hypertension as compared to AngII-treated WT mice (**Figure 7C**). These data are consistent with endothelial dysfunction and impaired vascular relaxation in eSWELL1 KO mice, resulting in a propensity for systolic hypertension.

**Figure 7.**
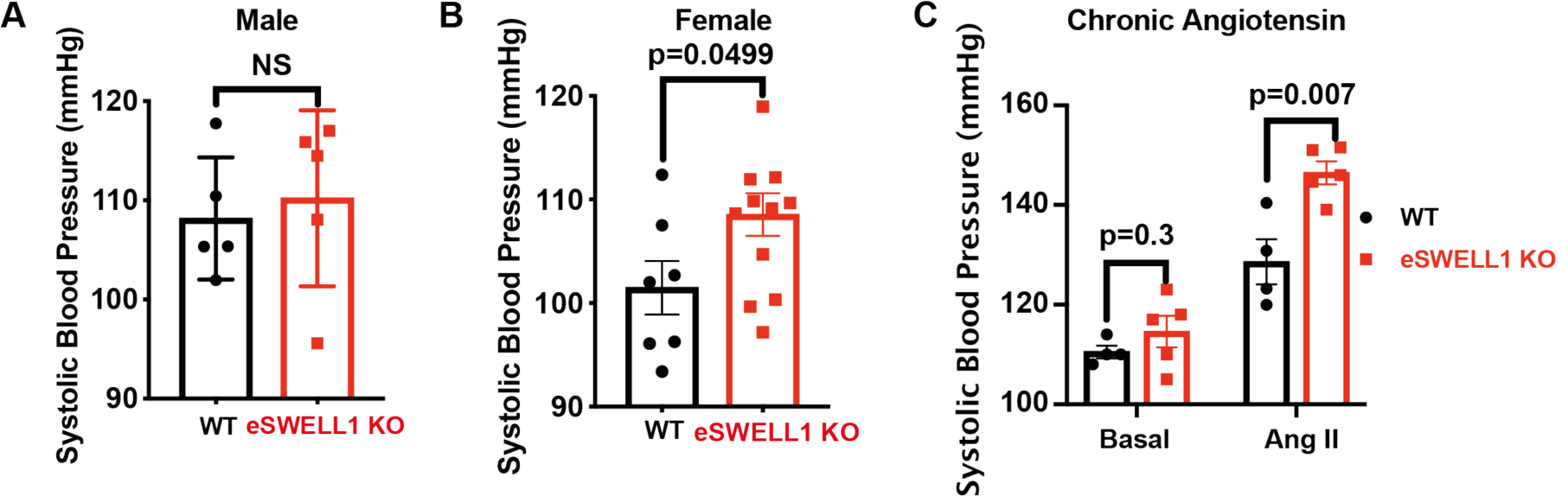
Endothelial-targeted SWELL1 deletion predisposes to systolic hypertension. Tail-cuff systolic blood pressures of (**A**) male and (**B**) female WT (n = 5 males and 7 females) and eSWELL1 KO (n = 5 males and 12 females) mice. (**C**) Systolic blood pressures of male WT (n = 4) and eSWELL1 KO (n = 5) mice under basal conditions and after 4 weeks of chronic angiotensin II infusion. Statistical significance between the indicated values calculated using a two-tailed Student’s t-test. P-values are illustrated on figures. Data are shown as mean ± s.e.m.

As endothelial dysfunction may also result in impaired blood flow we performed retinal imaging during i.p. injection of fluorescein to assess retinal vessel blood flow and morphology in WT and eSWELL1 KO mice. Mice raised on a regular diet have mild, non-significant impairments in retinal blood flow, based on the relative rate of rise of the fluorescein signal in retinal vessels (**Figure 8-Figure Supplement 1**). There is also evidence of mild focal narrowing of retinal vessels in eSWELL1 KO as compared to WT mice (**Figure 8-Figure Supplement 1A, D&E**), with no significant differences in other parameters (**Figure 8-Figure Supplement 1G-L**). In mice raised on high-fat high-sucrose (HFHS) diet, retinal blood flow is more severely impaired (**Figure 8A-C**) with significant focal, and diffuse retinal vessel narrowing in eSWELL1 KO mice compared to WT mice (**Figure 8A; F-H, Figure 8-Figure Supplement 2, Figure 8-Video 1**), and this relative difference is markedly worse in female compared to male mice. These findings are all consistent with endothelial dysfunction and impaired retinal vessel vasorelaxation due to reduced eNOS expression and activity, particularly in the setting of HFHS diet. Also consistent with impaired eNOS activity are reductions in vessel number (**Figure 8E**), vessel surface area (**Figure 8H**), number of end points (**Figure 8K**), branching index (**Figure 8L**), and increased lacunarity (**Figure 8J**). These parameters are all suggestive of diabetes-induced retinal vessel dysfunction in the eSWELL1 KO mice, consistent with the loss of eNOS activity that is expected when insulin signaling is compromised(48) (49). Notably, both WT and eSWELL1 KO mice were found to be equally glucose intolerant and insulin-resistant (**Figure 8-Figure Supplement 3**), indicating that these differences in microvascular dysfunction were not due increased hyperglycemia and more severe diabetes in eSWELL1 KO mice. Taken together, our findings reveal that SWELL1 is highly expressed in endothelium and functionally encodes endothelial VRAC. SWELL1 regulates ERK, AKT-eNOS, and mTOR signaling, forms a SWELL1-GRB2-Cav1-eNOS signaling complex, and regulates vascular function *in vivo*.

**Figure 8- Figure Supplement-1.**
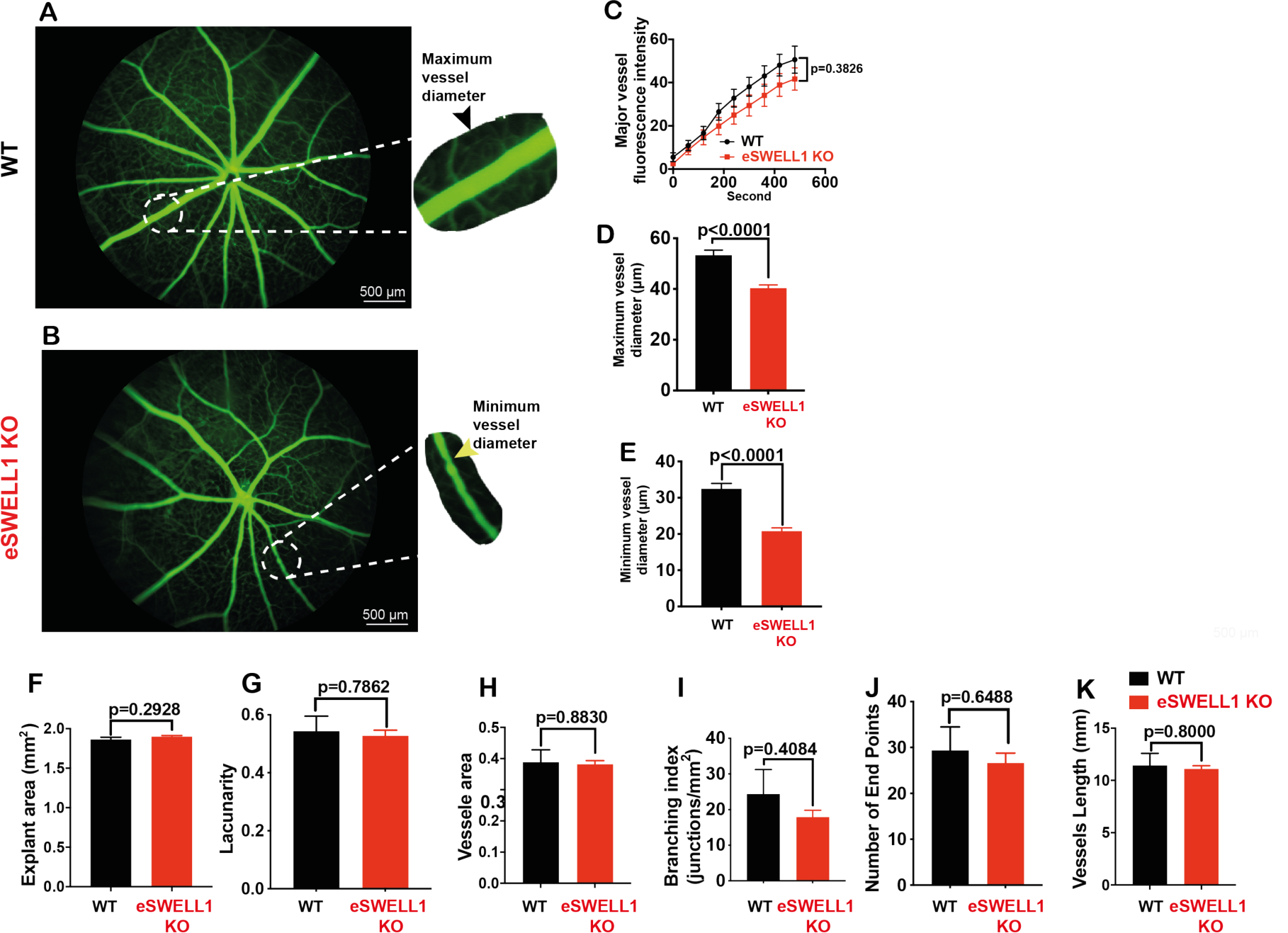
Endothelium-specific SWELL1 KO mice exhibit mild retinal microvascular disease at baseline. Representative fluorescein retinal angiograms of 13 week old WT (**A**) and eSWELL1 KO (**B**) mice raised on a regular diet. Inset shows magnified view of retinal vessels. (**C-E**) Quantification of major vessel fluorescence intensity over time after i.p. fluorescein injection (**C**), maximum vessel diameter (**D**), and minimum vessel diameter (**E**) in WT (n = 11 mice) and eSWELL1 KO (n = 11 mice). (**F-K**) Quantification of explant area (**F**), Lacunarity (**G**), Vessel area (**H**), Branching Index (**I**), Number of end points (**J**); and vessel length (**K**) in WT and eSWELL1 KO mice. Statistical significance between the indicated values calculated using a two-tailed Student’s t-test. P-values are illustrated on figures. Error bars represent mean ± s.e.m. Scale bar is 500 µm.

**Figure 8.**
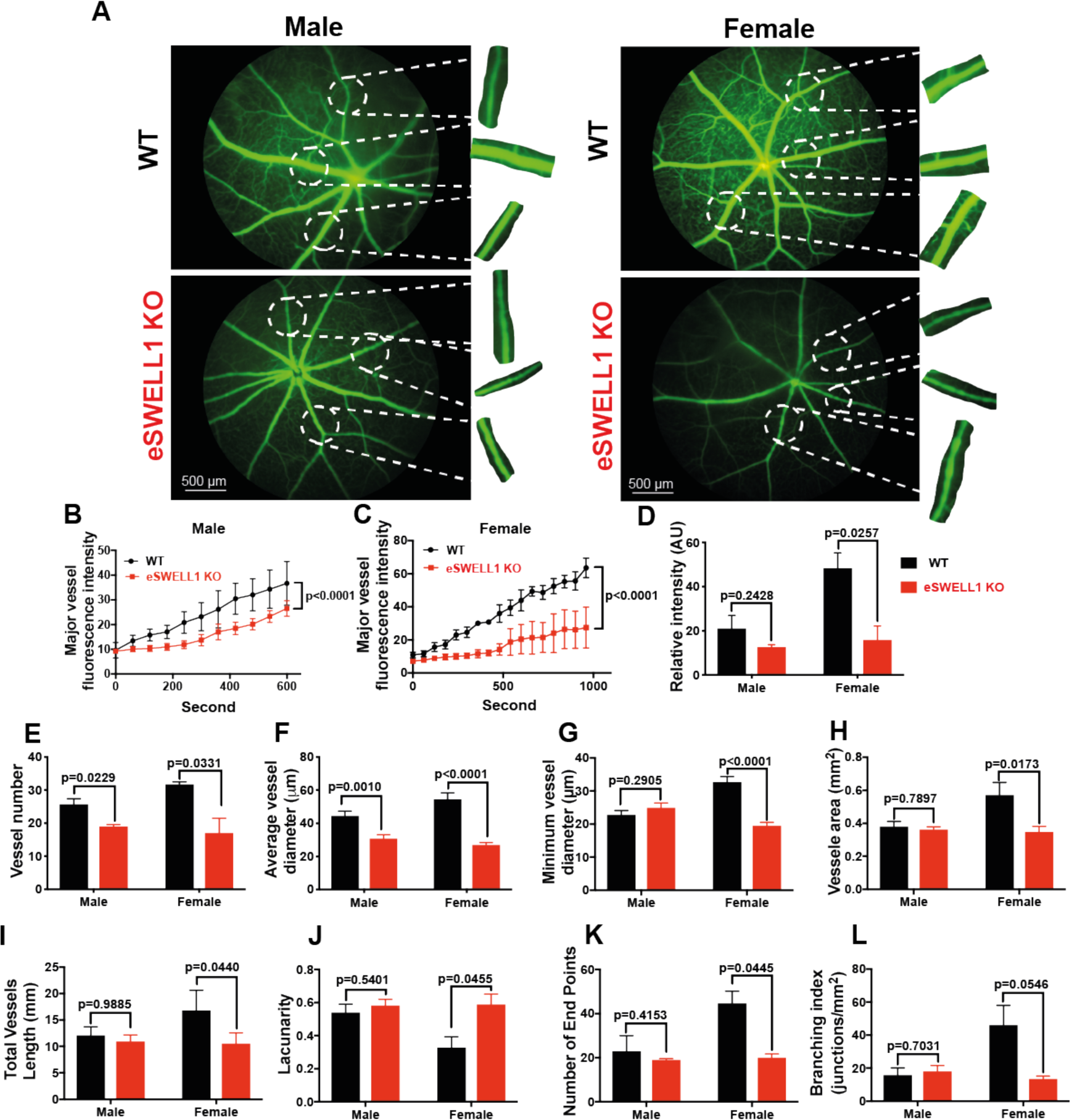
Endothelium-specific SWELL1 KO mice exhibit exacerbated impairments retinal microvascular disease in the setting of Type 2 diabetes. **A**, Representative fluorescein retinal angiograms of WT (top) and eSWELL1 KO (bottom) male (left) and female (right) mice raised on a high-fat high sucrose (HFHS) diet. Inset shows magnified view of retinal vessels. Quantification of major vessel fluorescence intensity over time after i.p. fluorescein injection in (**B**) male (n = 3) and (**C**) female (n = 3) WT and eSWELL1 KO mice. (**D-K**) Quantification of total retinal vessel intensity (**D**), Total vessel number (**E**); Vessel diameter (F); Minimum vessel diameter (**G**); Vessel area (**H**); Total vessel length (**I**); Lacunarity (**J**); Number of end points (**K**); and Branching index (**L**) of retinal vessels in WT and eSWELL1 KO mice. Statistical significance between the indicated values calculated using 2-way Anova for **B&C** and **D-L** using a two-tailed Student’s t-test. P-values are illustrated on figures. Error bars represent mean ± s.e.m.

**Figure 8- Figure Supplement-2.**
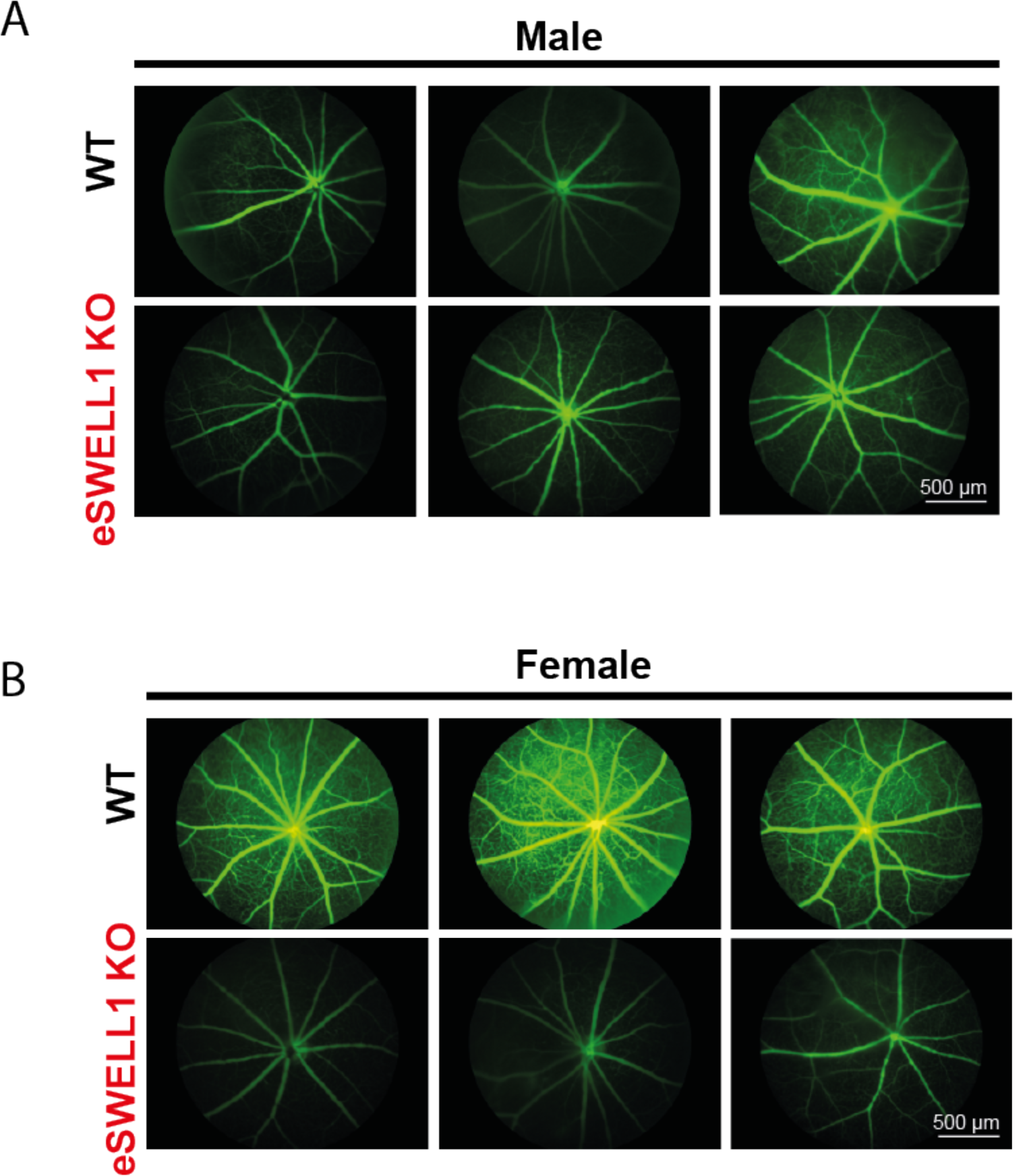
Endothelium-specific SWELL1 KO mice exhibit exacerbated impairments retinal microvascular disease in the setting of Type 2 diabetes. Three representative fluorescein retinal angiograms of WT and eSWELL1 KO from male (**A**) and female (**B**) mice raised on a high-fat high sucrose (HFHS) diet for 10 months. Scale bar is 500 µm.

**Figure 8- Figure Supplement-3.**
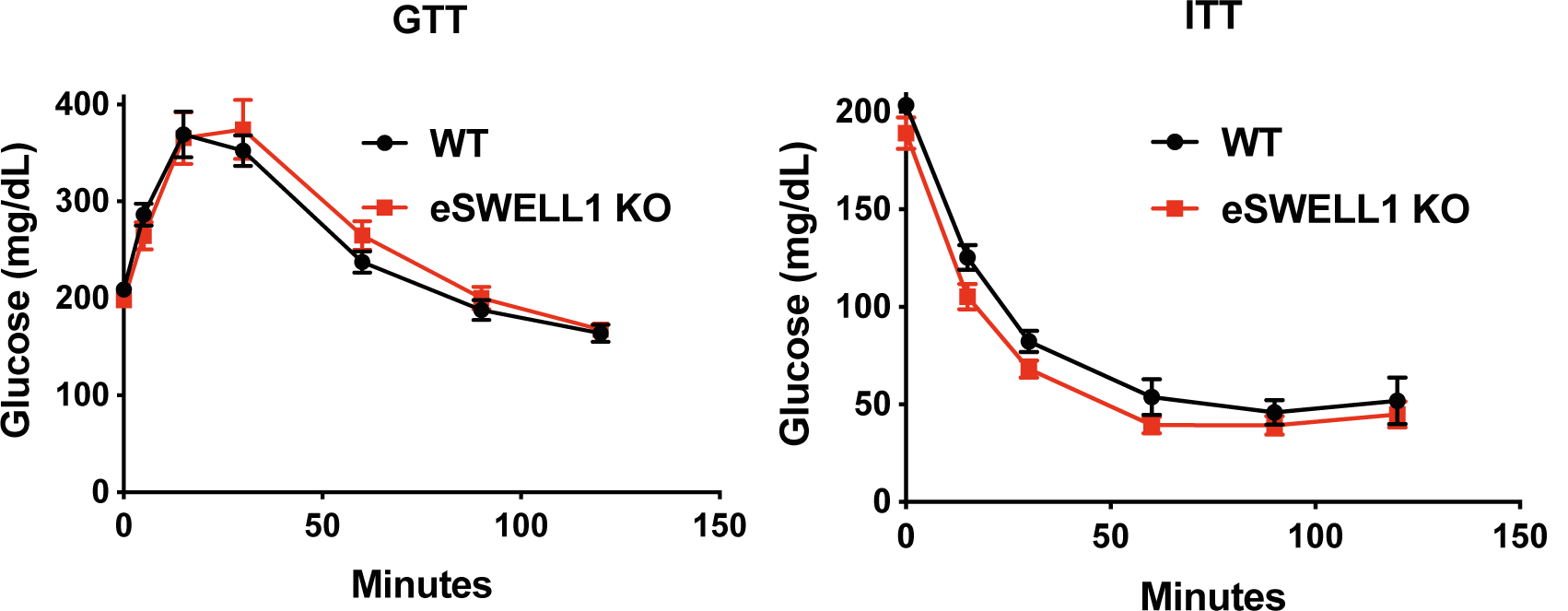
Glucose tolerance (GTT) and insulin tolerance (ITT) are not altered in endothelium-specific SWELL1 KO mice (n = 13) compared to WT mice (n = 8) raised on a high-fat high sucrose diet for 10 months.

## Discussion

Our findings demonstrate that the SWELL1-LRRC8 heterohexamer functionally encodes endothelial VRAC, whereby SWELL1-LRRC8 associates with GRB2 and Cav1 and positively regulates PI3K-AKT-eNOS and ERK1/2 signaling. Under basal conditions and with stretch, SWELL1 depletion in HUVECs reduces pAKT2, pAKT1, p-eNOS and pERK1/2. These data reveal that SWELL1 mediated PI3K-AKT signaling is conserved in endothelium, similar to previous observations in adipocytes(19), and in turn positively regulates eNOS expression and activity. Consistent with this mechanism, endothelial-targeted SWELL1 ablation *in vivo* predisposes to microvascular dysfunction in the setting of Type 2 diabetes, and to hypertension in response to angiotensin-II infusion. These results are in line with the notion that SWELL1 depleted endothelium contributes to an insulin-resistant state in which impaired PI3K-AKT-eNOS signaling results in a propensity for vascular dysfunction(24, 25). Insulin-mediated regulation of NO is physiologically(50-52) and pathophysiologically(53) important, as NO has vasodilatory(54, 55), anti-inflammatory(56), antioxidant(57), and antiplatelet effects(58-61). Indeed, impaired NO-mediated vascular reactivity is a predictor of future adverse cardiac events(62) and portends increased risk of atherosclerosis(63). Consistent with these NO-mediated effects, the RNA sequencing data derived from SWELL1 KD HUVECs revealed enrichment in inflammatory, cell adhesion, and proliferation pathways (GADD45, IL-8, mTOR, TREM1 signaling) that may arise from SWELL1-mediated dysregulation of eNOS activity.

In addition to reductions in AKT-eNOS signaling, SWELL1 depletion in HUVECs also reduced ERK1/2 signaling. This decrease in pERK1/2 suggests impaired MAPK signaling which is connected to the insulin receptor by GRB2-SOS. Indeed, we also found that SWELL1 and GRB2 interact in HUVECs, and this may provide the molecular mechanism for the observed defect in ERK signaling. Interestingly, GRB2-MAPK signaling is thought to promote angiogenesis, migration and proliferation(64), so reductions in ERK1/2 signaling would be predicted to inhibit these processes. Instead, aortic explants from SWELL1 KO mice exhibit augmented tube formation - indicative of pro-angiogenesis. This pro-angiogenesis, pro-migration cellular phenotype observed upon SWELL1 depletion might be explained by increases pS6K and p70 signaling, as observed in HUVECs, and suggestive of mTORC1 hyperactivation(47), in addition to the observed increases in VEGFA and CD31 expression. Future studies will further delineate the molecular mechanisms of SWELL1 modulation of insulin-GRB2-AKT/ERK1/2 and mTOR signaling in endothelial cells.

The phenotype of endothelial targeted SWELL1 KO (eSWELL1 KO) mice are consistent with a reduction in eNOS and p-eNOS as these mice exhibit mild hypertension at baseline (females) and exacerbated hypertension in response to chronic angiotension infusion, suggesting a modulatory effect on vascular reactivity. Similarly, retinal blood flow is only mildly impaired in eSWELL1 KO mice raised on a regular diet, with some evidence of microvascular disease. However, both retinal blood flow and retinal vessel morphology are markedly impaired in obese-T2D, insulin resistant eSWELL1 KO mice raised on a high fat high sucrose diet compared to controls. This is consistent with a synergistic role of endothelial SWELL1 ablation and T2D/obesity in the pathogenesis of vascular disease. Indeed, our results suggest that reductions in SWELL1 signaling may contribute to impaired vascular function observed in humans in response to insulin and/or shear stress in the setting of obesity (65-68) and insulin-resistance (69).

## AUTHOR CONTRIBUTIONS

Conceptualization, R.S.; Methodology, A.A., C.T., O.A., A.K., U.F., J.M., M. E-H., M.R., C.E.G., R.A.M., L.X., S.G., R.F.M., C.K., A.S., J.A., R.S.; Formal Analysis, A.A., R.S., C.T., J.M., A.K., L.X., S.G, R.A.M., U.F., C.K., C.E.G.; Investigation, R.S., A.A., L.X., J.M., A.K., S.G, C.T., C.K., C.G., R.M.; Resources, R.S.; Writing – Original Draft, R.S., Writing – Review & Editing, R.S., C.T., A.A., L.X., S.G., C.E.G.; Visualization, R.S., A.A., C.T., J.M., C.K., L.X., S.G, R.A.M., C.E.G.; Supervision, A.S., R.F.M., C.E.G, R.S.; Funding Acquisition, R.S.

## DATA AVAILABILITY

All raw RNA sequencing data will be uploaded to Gene Expression Omnibus (GEO). All other data will be made available upon reasonable request.

## ACKNOWLEDGMENTS

RNA-Seq data presented herein were obtained at the Genomics Division of the Iowa Institute of Human Genetics. This work was supported by grants from the NIH/NHLBI R01 HL125436 (C.E.G), NIH/NHLBI 5R00HL125683 (A.S), Cancer Research Foundation Young Investigator Award (A.S.), NIH NIDDK 1R01DK106009 (R.S.), the Roy J. Carver Trust (R.S.), UIHC Center for Hypertension Research Pilot and Feasibility Grant and from King Abdullah International Medical Research Center (KAIMRC) grant RA17-014-A (A. A.). We thank Dr. Rithwick Rajagopal for insightful reading of the manuscript.

## Materials and Methods

### Animals

The institutional animal care and use committee of the University of Iowa and Washington University School of Medicine approved all experimental procedures involving animals. All mice were housed in temperature, humidity and light controlled environment and allowed water access and food. Both male and female *Swell1*^*fl/fl*^ (1, 2)(WT), *CDH5*^*cre*^; *Swell1*^*fl/fl*^ (eSWELL1 KO) mice were generated and used in these studies. CHD5^Cre^ mice were obtained from Dr. Kaikobad Irani (University of Iowa, IA). In a subset of experiments, 5-8 week old *Swell1*^*fl/fl*^ and *CDH5* ^*cre*^; *Swell1*^*fl/fl*^ mice were switched to HFHS (High Fat high Sucrose rodent diet, Research Diets, Inc.,Cat # D12331) for at least 10 months. *CDH5*^*cre*^ mice were crossed with *Rosa26-tdTomato* (Jax# 007914) reporter mice to identify CDH5+ cells for primary endothelium patch-clamp studies.

### Antibodies

Rabbit polyclonal anti-SWELL1 antibody was generated against the epitope QRTKSRIEQGIVDRSE (Pacific Antibodies)(1). All other primary antibodies were purchased from Cells Signaling: anti-β-actin (#8457), Total Akt (#4685S), Akt1 (#2938), Akt2 (#3063), p-eNOS (#9571), Total eNOS (#32027), p-AS160 (#4288), p-p70 S6 Kinase (#9205S), pS6 Ribosomal (#5364S), GAPDH (#5174), pErk1/2 (#9101), Total Erk1/2 (#9102). Anti-SWELL1 antibody was custom made as described previously (1, 2). Purified mouse anti-Grb2 was purchased from BD (610111) and Santa Cruz (#sc-255). Rabbit IgG Santa Cruz (sc-2027). Anti-CD31 was purchased from Thermo Fisher (MA3105).

### Electrophysiology

All recordings were performed in the whole-cell configuration at room temperature, as previously described(1, 2). Briefly, currents were measured with either an Axopatch 200B amplifier or a MultiClamp 700B amplifier (Molecular Devices) paired to a Digidata 1550 digitizer. Both amplifiers used pClamp 10.4 software. The intracellular solution contained (in mM): 120 L-aspartic acid, 20 CsCl, 1 MgCl_2_, 5 EGTA, 10 HEPES, 5 MgATP, 120 CsOH, 0.1 GTP, pH 7.2 with CsOH. The extracellular solution for hypotonic stimulation contained (in mM): 90 NaCl, 2 CsCl, 1 MgCl_2_, 1 CaCl_2_, 10 HEPES, 5 glucose, 5 mannitol, pH 7.4 with NaOH (210 mOsm/kg). The isotonic extracellular solution contained the same composition as above except for mannitol concentration of 105 (300 mOsm/kg). The osmolarity was checked by a vapor pressure osmometer 5500 (Wescor). Currents were filtered at 10 kHz and sampled at 100 μs interval. The patch pipettes were pulled from borosilicate glass capillary tubes (WPI) by a P-87 micropipette puller (Sutter Instruments). The pipette resistance was ∼4-6 MΩ when the patch pipette was filled with intracellular solution. The holding potential was 0 mV. Voltage steps (500 ms) were elicited from 0 mV holding potential from −100 to +100 mV in 20 mV increments every 0.6 s. Voltage ramps from −100 to +100 mV (at 0.4 mV/ms) were applied every 4 s.

### Adenoviral knockdown

HUVECs were plated at 550,000 cell/well in 12 well plates. Cells were grown for 24 h in the plates and transduced with either human adenovirus type 5 with shLRRC8A (shSWELL1: Ad5-mCherry-U6-hLRRC8A-shRNA, 2.2×10^10^ PFU/ml, (shADV-214592), Vector Biolabs), or a scrambled non-targeting control (shSCR: Ad5-U6-scramble-mCherry, 1×10^10^ PFU/ml) at a multiplicity of infection (MOI) of 50 for 12 hours, and studies performed 3-4 days after adenoviral transduction. The shLRRC8a targeting sequence is: GCA CAA CAT CAA GTT CGA CGT.

### siRNA knockdown

HUVECs were plated at 360,000 cell/well in 6 well plate. Cells were grown for 24 h (90-95% confluency) and transduced with either a silencer select siRNA with si-LRRC8a (Cat#4392420, sense: GCAACUUCUGGUUCAAAUUTT antisense: AAUUUGAACCAGAAGUUGCTG, Invitrogen) or a non-targeting control silencer select siRNA (Cat# 4390846, Invitrogen), as described previously (3). The siLRRC8a used targets a different sequence from the shRNA described above. Briefly, siRNAs were transduced twice, 24 and 72 hours after HUVEC plating. Each siRNA was combined with Opti-MEM (285.25 µl, Cat#11058-021, Invitrogen) siPORT™ amine (8.75 µl, Cat#AM4503, Invitrogen) and the silencer select siRNA (6 µl) in a final volume of 300 µl. HUVECs were transduced over a 4-hour period at 37°C, using DMEM media +1% FBS. After transduction, the cells were returned to media containing M199, 20% FBS, 0.05 g Heparin Sodium Salt, and 15 mg ECGS. Cell lysates were collected at basal conditions on day 4.

### Isolation of mouse lung endothelial cells

Isolation of mouse lung endothelial cells was performed according to the following protocol: Day 1-Incubate sheep anti-rat IgG Dynabeads (Invitrogen) overnight with PECAM (Sigma, #SAB4502167) and VEGFR2 (R&D Systems, #BAF357) antibodies at 4 °C in PBS with gentle agitation. Day 2-lungs were removed from the mice, washed in 10% FBS/DMEM, minced into 1-2 mm squares and digested with Collagenase Type I (2 mg/ml, Gibco) at 37°C for 1 hour with agitation. The cellular digest was filtered through a 70 µm cell strainer, centrifuged at 1500 rpm and the cells immediately incubated with the antibody coated Dynabeads at room temperature for 20 minutes. The bead-bound cells were recovered with a magnet, washed two times with PBS, and plated overnight on collagen type I (100ug/ml) coated cover slips. The endothelial cells were maintained in a growth media of M199, 20% FBS, 0.05g Heparin Sodium Salt, 50mg/ml ECGS, and 1x Anti-Anti.

### Cells Culture

HUVECs were purchased from ATCC and were grown in MCDB-131-Complete media overnight. HUVECs for basal condition collection were grown in growth media of M199, 20% FBS, 0.05g Heparin Sodium Salt (Cat#9041-08-1, Alfa Aesar), and 15 mg ECGS (Cat#02-102, Millipore Sigma). Cells were routinely cultured on 1% of gelatin coated plates at 37°C at 5% CO_2_. For insulin stimulation (Cat#SLBW8931), cells were serum starved for at least 13 h in 1% FBS (Atlanta Bio selected, Cat #S11110) or without FBS using endothelial cells growth basal medium (Lonza cat#cc-3121) instead of MCDB-131-complete media. Insulin stimulation was used for the times indicated at 100 nM.

### Immunoblotting

Cells were harvested in ice-cold lysis buffer (150 mM NaCl, 20 mM HEPES, 1% NP-40, 5mM EDTA, pH 7.5) with added proteinase/phosphatase inhibitor (Roche). Cells were kept on ice with gentle agitation for 20 minutes to allow complete lysis. Lysate scraped into 1.5 ml tubes and cleared of debris by centrifugation at 14,000 x g for 20 minutes at 4 °C. Supernatant were transferred to fresh tube and solubilized protein was measured using a DC protein assay kit (Bio-Rad). For immunoblotting an appropriate volume of 1 x Laemmli (Bio-rad) sample loading buffer was added to the sample (10 μg of protein), which then heated at 90°C for 5 min before loading onto 4-20% gel (Bio-Rad). Proteins were separated using running buffer (Bio-Rad) for 2 h at 150 V. Proteins were transferred to PVDF membrane (Bio-Rad) and membrane blocked in 5% (w/v) BSA in TBST or 5 % (w/v) milk in TBST at room temperature for 2 hours. Blots were incubated with primary antibodies at 4 °C overnight, followed by secondary antibody (Bio-Rad, Goat-anti-mouse #170-5047, Goat-anti-rabbit #170-6515, all used at 1:10000) at room temperature for one hour. Membranes were washed3 times and incubated in enhanced substrate Clarity (Bio-Rad) and imaged using a ChemiDoc XRS using Image Lab (Bio-Rad) for imaging and analyzing protein band intensities. ß-Actin or GAPDH levels were quantified to correct for protein loading.

### Immunoprecipitation

Cells were seeded on gelatin-coated 10 cm dishes in complete media for 24 h. Adenoviruses, Ad5-mCherry-U6-hLRRC8A-shRNA or Ad5-U6-scramble-mCherry were added to cells for 12h. After 4 days cells were serum starved for 16 h with basal media contain 1% serum before stimulation with insulin (10 nM/ml). Cells were harvested in ice-cold lysis buffer (150 mM NaCl, 20 mM HEPES, 1% NP-40, 5mM EDTA, pH 7.5) with added proteinase/phosphatase inhibitor (Roche) and kept on ice with gentle agitation for 15 minutes to allow complete lysis. Lysated were incubated with anti-Grb2 antibody (20 μg/ml) or control rabbit IgG (20 μg/ml, Santa Cruz sc-2027) rotating end over end overnight at 4 °C. Protein G sepharose beads (GE) were added to this for a further 4 h before samples were centrifuged at 10,000 x g for 3 minutes and washed three times with RIPA buffer and re-suspended in laemmli buffer (Bio-Rad), boiled for 5 minutes, separated by SDS-PAGE gel followed by the western blot protocol.

### Stretch assay

Equal amounts of cells were plated in each well in 6 well plated BioFlex coated with Laminin (BF-3001CCase) culture plate and seeded to approximately 90% confluence. Plates were placed into a Flexcell Jr. Tension System (FX-6000T), and incubated at 37°C with 5% CO_2_. Prior to stretch-stimulation, basal media of 1% FBS was added for 16 h. Cell on flexible membrane were subjected to static stretch with following parameters: a stretch of 0.1% and 5% with static strain. Cells were stretched for 5, 30, 60 or 180 minutes. Cells were then lysed and protein isolated for subsequent Western blots.

### Immunofluorescence imaging

Cells were plated on gelatin-coated glass coverslips. Cells on cover slip were washed in PBS and fixed with 2% (w/v) PFA for 20 minutes at room temperature. PFA were washed three times with PBS and permeabilized in PBS containing 0.2% Triton X-100 for 5 minutes at room temperature. Cells on coverslips were washed in PBS and blocked for 30 minutes at room temperature with TBS containing 0.1% Tween-20 and 5% BSA. Cells on coverslip were incubated overnight at 4°C with primary antibody (1:250) in TBS containing 0.1% Tween-20 and 1% BSA. Cells were then washed in PBS 5x and incubated for 2 hours at room temperature with 5% BSA in TBST. Cells were washed three times in TBST and then incubated with secondary antibodies at 1:1000 dilution (Invitrogen, Anti-Rabbit 488, A11070; Anti-mouse 568, A11019; Anti-mouse 488, A11017) for 1 h at room temperature. Coverslips were then incubated for 10 minutes with Topro 3 (T3605, Thermofisher) or mounted with mounting media containing DAPI (Invitrogen), to visualize nuclei. Images were taken using Axiocam 503 Mono Camera controlled by Zeiss Blue using a Plan-Apochromate 40x oil immersion objective.

### Ex vivo sprouting angiogenesis assay

Following Avertin injection and cervical dislocation, aortas were dissected and connective tissue removed, and then washed with PBS with 50 μg/ml penicillin and streptomycin. Using iris scissors, the aorta was cut into aortic rings of 1∼2 mm cross sectional slices. 50 μl of Matrigel was used to coat the center of coverslips in 24 well plates for two hours at 37 °C in the incubator to solidify the Matrigel. Aorta rings were then seeded and transplanted on Matrigel (BD Biosciences, Cat#356231) on coverslips. After seeding the aortic rings, plates were incubated in 37 °C without medium for 10 minutes to allow the ring to attach to the Matrigel. Complete medium was added to each well and incubated at 37°C with 5% CO_2_ for 48-72 h. Phase contrast photos of individual explants were taken using a 10x/0.75 NA objective Olympus IX73 microscope (Olympus, Japan) fitted with camera (Orca flash 4.0+, Hamamatsu, Japan). The areas of sprouting, number of tips cells and length of the tube for each condition were quantified with computer software ImageJ 1.52i (National Institute of Health). Cells were incubated with SWELL1 (1:250) and CD31 (1:250) primary antibodies in 0.1% Tween-20 and 1% BSA overnight, and then incubated with secondary antibodies (Invitrogen, Anti-Rabbit 488, A11070; Anti-Mouse 568, A11019). Cells were then incubated for 10 minutes with Topro 3 (T3605, Thermofisher) at room temperature.

### Retina imaging

Ketamine (Akorn Animal Health, 100mg/ml) was prepared and mixed with Xylazine then stored at room temperature. Animals were anesthetized with 87.5/12.mg/kg BW via intraperitoneal (IP) injection. Eyes were topically anesthetized with proparacaine and dilated with tropicamide. Fluorescein (100 mg/ml, Akorn Inc) diluted with sterile saline was administered by IP injection (50 μl), mice positioned on Micron imaging platform. Images of the eyes were taken from the start of fluorescein infusion with the Micron camera with a 450-650 nm excitation filter and 469-488 nm barrier filter at 30 frame/sec using Micron software for 30 seconds. Data were converted into tiff image for further analysis.

### Angiotensin-II infusion

Infusion studies carried out using Azlet osmotic minipumps (Model 1004). Angiotensin-II (BACHEM) dissolved in saline was filled in the minipumps and were prepared to maintain infusion rate of 600 ng/kg/min for four weeks. The mice were anesthetized under 2% isofluorane and the minipumps were implanted subcutaneously on the dorsal aspect of the mice.

### Blood pressure recordings

Systolic tail-cuff blood pressure (BP) measurements were carried out using computerized tail-cuff system BP-2000 (Visitech Systems) at the same time of day. Mice were first acclimated to the device by performing 3 days of measurements (20 sequential measurements/day) and then mean blood pressure readings were obtained by averaging 3-5 days of measurements (not inclusive of the 3 acclimation days).

### RNA sequencing

RNA quality was assessed by Agilent BioAnalyzer 2100 by the University of Iowa Institute of Human Genetics, Genomics Division. RNA integrity numbers greater than 8 were accepted for RNAseq library preparation. RNA libraries of 150 bp PolyA-enriched RNA were generated, and sequencing was performed on a HiSeq 4000 genome sequencing platform (Illumina). Sequencing results were uploaded and analyzed with BaseSpace (Illumina). Sequences were trimmed to 125 bp using FASTQ Toolkit (Version 2.2.0) and aligned to Mus musculus mmp10 genome using RNA-Seq Alignment (Version 1.1.0). Transcripts were assembled and differential gene expression was determined using Cufflinks Assembly and DE (Version 2.1.0). Ingenuity Pathway Analysis (QIAGEN) was used to analyze significantly regulated genes which were filtered using cutoffs of >1.5 fragments per kilobase per million reads, >1.5 fold changes in gene expression, and a false discovery rate of <0.05. Heatmaps were generated to visualize significantly regulated genes. Data have been deposited in GEO (accession# TBD).

### Metabolic phenotyping

Mice were fasted for 6 h prior to glucose tolerance tests (GTT). Baseline glucose levels at 0 min timepoint (fasting glucose, FG) were measured from blood sample collected from tail snipping using glucometer (Bayer Healthcare LLC). 0.75 g D-Glucose/kg body weight were injected (i.p.) for HFHS mice, respectively and glucose levels were measured at 7, 15, 30, 60, 90 and 120 min timepoints after injection. For insulin tolerance tests (ITTs), the mice were fasted for 4 h. Similar to GTTs, the baseline blood glucose levels were measured at 0 min timepoint and 15, 30, 60, 90 and 120 min timepoints post-injection (i.p.) of insulin (HumulinR, 1.25 U/kg body weight).

